# A spatiotemporal translatome of mouse tissue development

**DOI:** 10.1101/2020.04.14.041079

**Authors:** Hongwei Wang, Yan Wang, Jiaqi Yang, Nan Tang, Huihui Li, Mingzhe Xie, Zhi Xie

## Abstract

The precise regulation of gene expression in mammalian tissues during development results in their functional specification. Although previous transcriptomic and proteomic analyses have provided great biological insights into tissue-specific gene expression and the physiological relevance of these tissues in development, our understanding of translational regulation in developing tissues is lacking. In this study, we performed a spatiotemporally resolved translatome analysis of six mouse tissues at the embryonic and adult stages to quantify the effects of translational regulation and identify new translational components. We quantified the spatial and temporal divergences of gene expression and detected specific changes in gene expression and pathways underlying these divergences. We further showed that dynamic translational control can be achieved by modulating the translational efficiency, which resulted in the enhancement of tissue specificity during development. We also discovered thousands of actively translated upstream open reading frames (ORFs) that exhibited spatiotemporal patterns and demonstrated their regulatory roles in translational regulation. Furthermore, we identified known and novel micropeptides encoded by small ORFs from long noncoding RNAs that are functionally relevant to tissue development. Our data and analyses facilitate a better understanding of the complexity of translational regulation across tissue and developmental spectra and serve as a useful resource of the mouse translatome.

## Introduction

Mammalian tissues show extreme functional diversity despite sharing almost identical genetic codes. Their unique physiological functions are achieved through precise orchestration of spatiotemporal changes in gene expression during development. The quantification of gene expression across diverse tissues and developmental stages is vital for understanding the molecular and mechanistic principles underlying morphogenesis. Transcriptomic studies have characterized the gene expression profile in a variety of mammalian tissues during development and have thus revealed the complexity and dynamics of the transcriptome (Cardoso-Moreira et al. 2019; Sarropoulos et al. 2019). Proteomic studies have resolved the molecular details of the proteome variations in different mammalian tissues and thus extended our understanding of the spatiotemporal programs of protein expression (Uhlen et al. 2015; Zhou et al. 2017).

Transcription and translation are the two major steps in gene expression. During translation, ribosomes perform protein synthesis to ensure that a genetic code is successfully translated into a functional protein (Sonenberg and Hinnebusch 2009). Although transcriptomic and proteomic analyses have provided great biological insights into tissue specificity and the physiological relevance of tissues in development, the gene translation profiles in developing tissues have not been systematically investigated. Ribosome profiling (Ribo-seq) enables genome-wide quantitative measurements of gene translation at nucleotide resolution (Ingolia et al. 2009). By pinpointing ribosomes during translation, this technique allows a detailed analysis of the ribosome density on individual RNAs, the identification of canonical translated open reading frames (ORFs) and the discovery of noncanonical translated upstream ORFs (uORFs) in 5’UTRs and small ORFs (smORFs) in noncoding RNAs (Ingolia 2014; Brar and Weissman 2015; Ingolia 2016; Ingolia et al. 2019). Thus, a multi-tissue and multi-developmental-stage survey of the translational landscape will provide insight into key translational components and regulation processes underlying the physiology of tissue specificity and development.

This study constitutes the first spatiotemporally resolved translatome analysis of mouse tissues during development. We analyzed six mouse tissues at the embryonic and adult stages to quantify the effects of translational regulation and identify new translational components. We quantified the spatiotemporal divergences in gene expression and detected specific changes in gene expression and functions underlying these divergences. We further showed that dynamic translational control can be achieved by modulating the translational efficiency, which results in enhanced tissue specificity during development. We also discovered thousands of uORFs that exhibit spatiotemporal patterns and demonstrated their regulatory roles in translational regulation. Furthermore, we identified micropeptides encoded by smORFs from long noncoding RNAs that are functionally relevant to tissue development. Our data and analyses facilitate a better understanding of the complexity of translational regulation across tissue and developmental spectra and can serve as an important resource of the mouse translatome. The data and analyses are publicly available at http://sysbio.gzzoc.com/MSTA/.

## Results

### Transcriptional and translational maps of mouse tissue development

To obtain a global view of gene expression during the development of mammalian tissues, we performed Ribo-seq and RNA sequencing (RNA-seq) to profile six tissues from wild-type C57BL/6 mice at embryonic day (E) 15.5 and postnatal day (P) 42 (**Fig. 1a**). These tissues included ectoderm-derived brain and retinal tissues, mesoderm-derived heart and kidney tissues, and endoderm-derived liver and lung tissues. In total, the Ribo-seq experiments yielded more than 2.58 billion raw reads, with an average of ~107 million reads per library, and the RNA-seq experiments yielded more than 1.19 billion raw reads, with an average of ~50 million reads per library (**Table S1**). The ribosome-protected footprints (RPFs) obtained from the Ribo-seq analyses showed a predominant length of 29-30 nucleotides (**Fig. 1b**) with a strong preference for 5’untranslated regions (UTRs) compared with RNA-seq reads (**Fig. S1a**). On average, 76.4% of the RPFs were mapped to annotated coding sequences (CDSs), whereas 10.2% were mapped to 3’UTRs, 7.3% were mapped to intronic sequences (introns), and 6.1% were mapped to 5’UTRs. A reading frame analysis of the RPFs mapped to annotated CDS regions revealed a clear three-nucleotide (3-nt) periodicity corresponding to codon triplets, and 83% of the RPFs accumulated in the first frame, which indicated that these displayed a clear frame preference (**Fig. 1c-d**). As expected, the RNA-seq data did not show 3-nt periodicity or frame preference. Our experiments were highly reproducible, as indicated by nearly perfect correlations among the biological replicates (mean Pearson correlation coefficients r=0.9878 and 0.9954 for Ribo-seq and RNA-seq, respectively) (**Fig. S1b**). A principal component analysis (PCA) showed distinct separation of both the Ribo-seq and RNA-seq data among different stages and tissues (**Fig. S1c**). Taken together, these results demonstrate that our sequencing data are of high quality.

**Figure 1.**
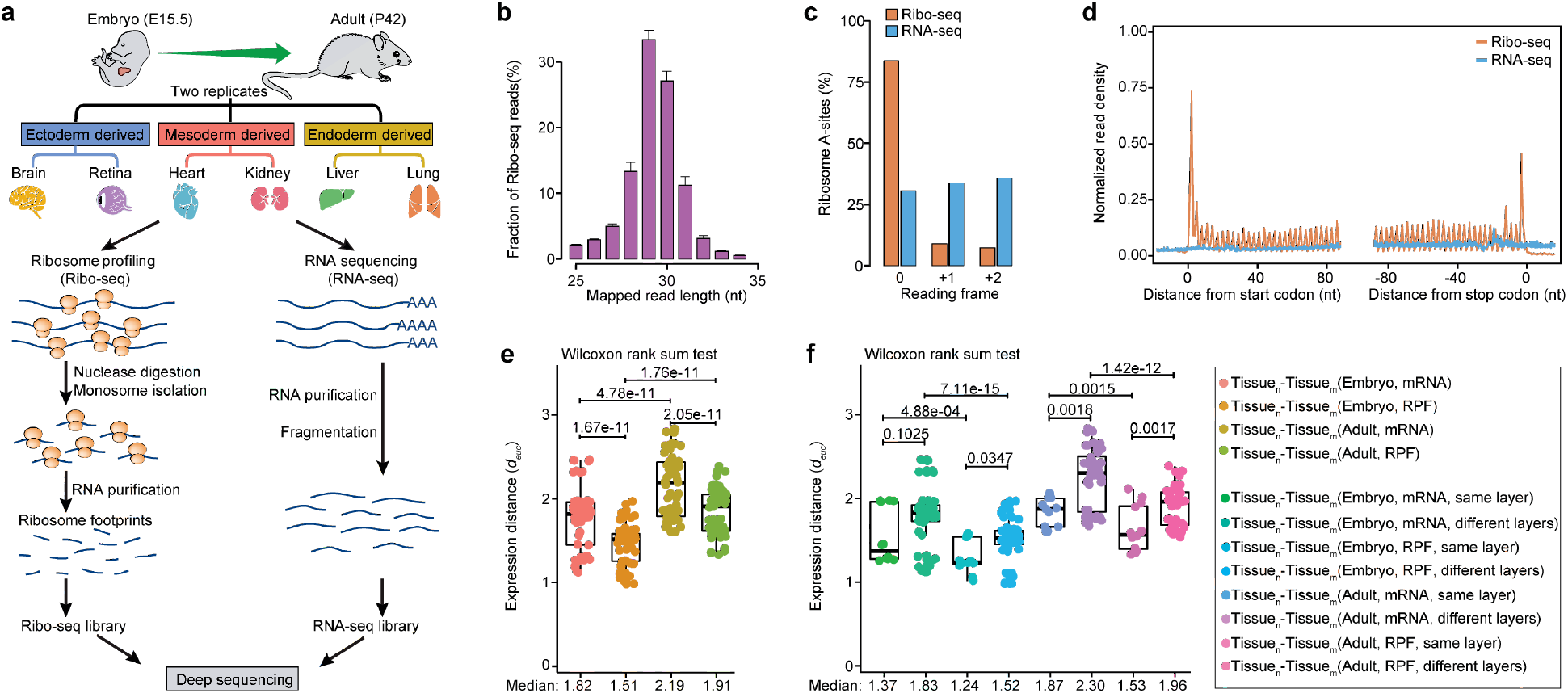
Patterns of gene expression during tissue development. (**a**) Brief overview of the experimental design. Detailed step-by-step protocols for the Ribo-seq and RNA-seq experiments can be found in the **Methods**. (**b**) Size distribution of the mapped Ribo-seq reads, with peaks at 29-30 nt. The error bars represent the standard errors of the mean (SEs). (**c**) Reading frame analysis of uniquely mapped Ribo-seq reads and RNA-seq reads within the annotated CDS regions. (**d**) Distribution of uniquely mapped reads along the CDS within each codon. Each read was reduced to a specific A-site nucleotide position depending on its fragment length. (**e**) Differences in the global gene expression profiles among different expression layers and developmental stages. (**f**) Differences in global gene expression profiles among germ layer-derived tissues. Each point represents the divergence in gene expression profiles between a pair of tissues. The between-group differences were compared using a Wilcoxon rank sum test, and the *P* values are shown. The boxplots show the medians, first quartiles and third quartiles; the lines extend to the furthest value within 1.5 of the interquartile range, and the gray points represent the mean values.

### Spatiotemporal divergence of gene expression during tissue development

Using the generated datasets, we investigated the global changes in gene expression among different expression levels, tissues, and developmental stages. We measured the divergence between a pair of expression profiles based on the Euclidean distance (Glazko and Mushegian 2010) (see **Methods**). We first compared the gene expression divergences in the six tissues between transcriptional and translational levels. For both the embryonic and adult stages, the inter-tissue divergence at the translational level was significantly smaller than that at the transcriptional level (**Fig. 1e**). For the embryonic and adult stages, the median expression distances among tissues at the translational level relative to that at the transcriptional level were 83% and 87%, respectively, which suggests that post-transcriptional gene regulation, particularly translational control, is responsible for buffering the inter-tissue expression divergence in transcription. As expected, the transcriptional and the translational divergences between tissues derived from the same germ layer were consistently smaller than those between pairs of tissues derived from different germ layers at both the embryonic and adult stages (**Fig. 1f**), which suggests that tissues from the same germ layer globally exhibit similar gene expression.

We subsequently compared the divergences in gene expression of the six tissues between the different developmental stages. Interestingly, at both the transcriptional and translational levels, the inter-tissue divergence at the adult stage was significantly larger than that at the embryonic stage, which suggests the gene expression landscape of tissues exhibits a higher diversity in adults than in embryos (**Fig. 1e**). The median expression distance between tissues at the adult stage relative to that at the embryonic stage exhibited a 1.2-fold increase at the transcriptional level and an even higher increase of 1.3-fold at the translational level. These trends in global expression patterns among different expression levels and developmental stages were consistent with the results from the tissue-specificity analysis (**Fig. S2**). Taken together, our analyses quantify the different degrees of gene expression divergences among different expression levels, tissues, and developmental stages and indicate that gene expression is potentially regulated during tissue development.

### Analysis of genes and pathways underlying gene expression divergence

We then sought to characterize the genes and pathways underlying the detected divergences in gene expression. Based on the transcriptional or translational expression levels, we classified all the protein-coding genes into five major categories, namely, ‘tissue-enriched’, ‘group-enriched’, ‘expressed-in-all’, ‘mixed’, and ‘not-expressed’ (**Fig. 2a** and **Table S2**; see **Methods**). The five classes showed distinct divergences in gene expression, particularly between the tissue-enriched and expressed-in-all classes (**Fig. 2b**). A Gene Ontology (GO) enrichment analysis revealed functional differences underlying the expression divergences. In particular, the tissue-enriched class showed large expression divergence and exhibited functional specificity, and the enriched GO terms coincided well with the main physiological roles of the respective tissue (**Table S3**). The comparison of the enriched GO terms obtained for each tissue at the translational level with those at the transcriptional level showed that the translational level contributed an average of 26% specific terms (**Fig. 2c**). For instance, some GO terms related to kidney function, such as ‘bile acid and bile salt transport’, ‘drug transport’, and ‘calcium ion homeostasis’, were enriched in kidney-specific translated genes, whereas some GO terms related to liver function, such as ‘bile acid biosynthetic process’, ‘lipid metabolic process’, and ‘vitamin transport’, were enriched in liver-specific translated genes. Moreover, some immunological functions, such as ‘regulation of innate immune response’ and ‘positive regulation of T cell proliferation’, were also found in translation-specific GO terms, and this finding could be explained by the fact that the liver forms part of the mononuclear phagocyte system and contains a high number of Kupffer cells involved in immune activity. These results illustrate the importance of the tissue-specific translatome in the maintenance of tissue functionality.

**Figure 2.**
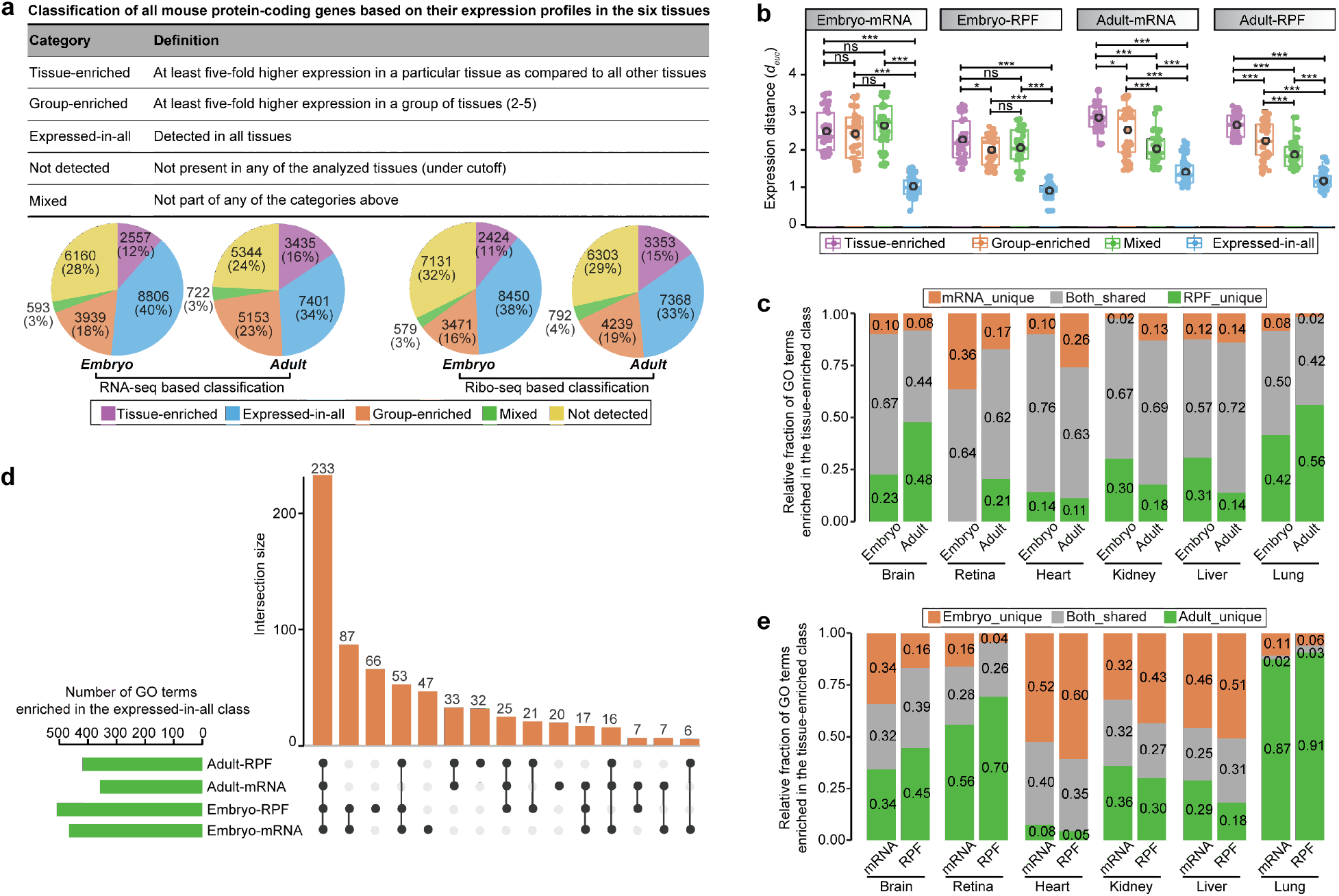
Genes and pathways underlying gene expression divergences. (**a**) Classification of mouse protein-coding genes based on their expression levels in the six tissues. The pie charts show the numbers and percentages of genes in each class. (**b**) Differences in global gene expression profiles among different classes of genes. The between-group differences were compared using a Wilcoxon rank sum test, and the *P* values are shown. ***: *P*<0.001; **: *P*<0.01; *: *P*<0.05; and ns: not significant. (**c**) Relative fractions of shared and unique GO terms enriched in the tissue-enriched class for each tissue between the transcriptional and translational levels. (**d**) Numbers of enriched GO terms in the expressed-in-all class and the overlap of enriched GO terms across different expression levels and developmental stages. (**e**) Relative fractions of shared and unique GO terms enriched in the tissue-enriched class for each tissue between the embryonic and adult stages.

In contrast to the tissue-enriched class, the expressed-in-all class showed a small divergence in expression and exhibited functional generality, which is generally found for the functional terms of housekeeping genes, such as those required for the maintenance of basal cellular functions involving many essential steps of gene expression, ranging from transcription initiation to RNA processing and post-translational modification (**Table S3**). Notably, an uncoupling of functional changes between transcription and translation was also found, and an average of 20% GO terms, including ‘ribosome disassembly’, ‘miRNA mediated inhibition of translation’, ‘IRES-dependent viral translational initiation’, and ‘posttranscriptional regulation of gene expression’, were uniquely enriched at the translational level (**Fig. 2d**). These results highlight the significance of translation in shaping the physiological functions necessary for all tissues.

We further analyzed the changes underlying the gene expression divergences observed during development. Along with tissue development and maturation, the fraction of tissue-enriched genes gradually increased, but the fraction of expressed-in-all genes gradually decreased (**Fig. 2a)**. Coinciding with this transition in gene expression patterns, the enriched GO terms obtained for each tissue also showed remarkable developmental stage specificity (**Fig. 2d-e** and **Table S3**). For instance, the retina-specific genes of adult mice were greatly involved in light stimulus-related functions, such as ‘detection of light stimulus involved in visual perception’, ‘sensory perception of light stimulus’, and ‘cellular response to light stimulus’. These functions are related to enhancing visual functions, which might be a result of increases in visual stimuli and interactions with light that promote neural plasticity in the retina after eye opening. The lung-specific genes of adult mice were more significantly involved in immunity-related functions compared with their embryonic counterparts, which is likely a manifestation of immune development and maturation during the postnatal period. Taken together, these results highlight the specific role that translation plays in tissue development and provide novel biological insights into genes and pathways that are important for tissue functionality.

### Coordination of transcriptional and translational regulation during development

We subsequently sought to understand how transcriptional and translational regulation coordinatively contribute to the divergences in the spatiotemporal patterns of gene expression. We first performed differential transcription and translation analyses among different tissues and developmental stages, which yielded large numbers of tissue- and developmental stage-specific differentially expressed genes (**Fig. 3a-b** and **Fig. S3**). A graph-based clustering analysis was subsequently applied using the similarity matrix of all differentially transcribed and translated genes across each pairwise comparison to delineate different gene clusters that are expected to show co-regulatory functional arrangements. Each comparison identified an average of 31 (range 21-37) potential clusters of co-regulated genes (**Fig. 3c** and **Fig. S4a**). To dissect the relative contribution of transcriptional and translational regulation to the overall differential patterns of each gene cluster, we performed a PCA (see **Methods**), which revealed the manifestations of individual clusters within the global regulatory programs (**Fig. 3d** and **Fig. S4b**). Some gene clusters were subjected to major transcriptional or translational regulation, whereas others were subjected to combinatorial regulation. A GO enrichment analysis of the gene clusters further showed their specific regulatory functions (**Table S4**). Taking the brain as an example, cluster 23 was primarily under translational regulation and enriched in translation-related functions, such as ‘cytoplasmic translation’, ‘translational initiation’, and ‘translational elongation’ (**Fig. 3e**), which facilitate immediate cellular adjustment through direct regulation of protein synthesis rather than transcriptional regulation. Cluster 8 was under both transcriptional and translational regulation and enriched in cellular signaling-related functions, such as ‘signal transduction’, ‘G protein-coupled receptor signaling pathway’, and ‘neuropeptide signaling pathway’, which reflect signaling-mediated developmental effects. Taken together, these results reveal that diverse regulatory controls of transcription and translation contribute to tissue development and function.

**Figure 3.**
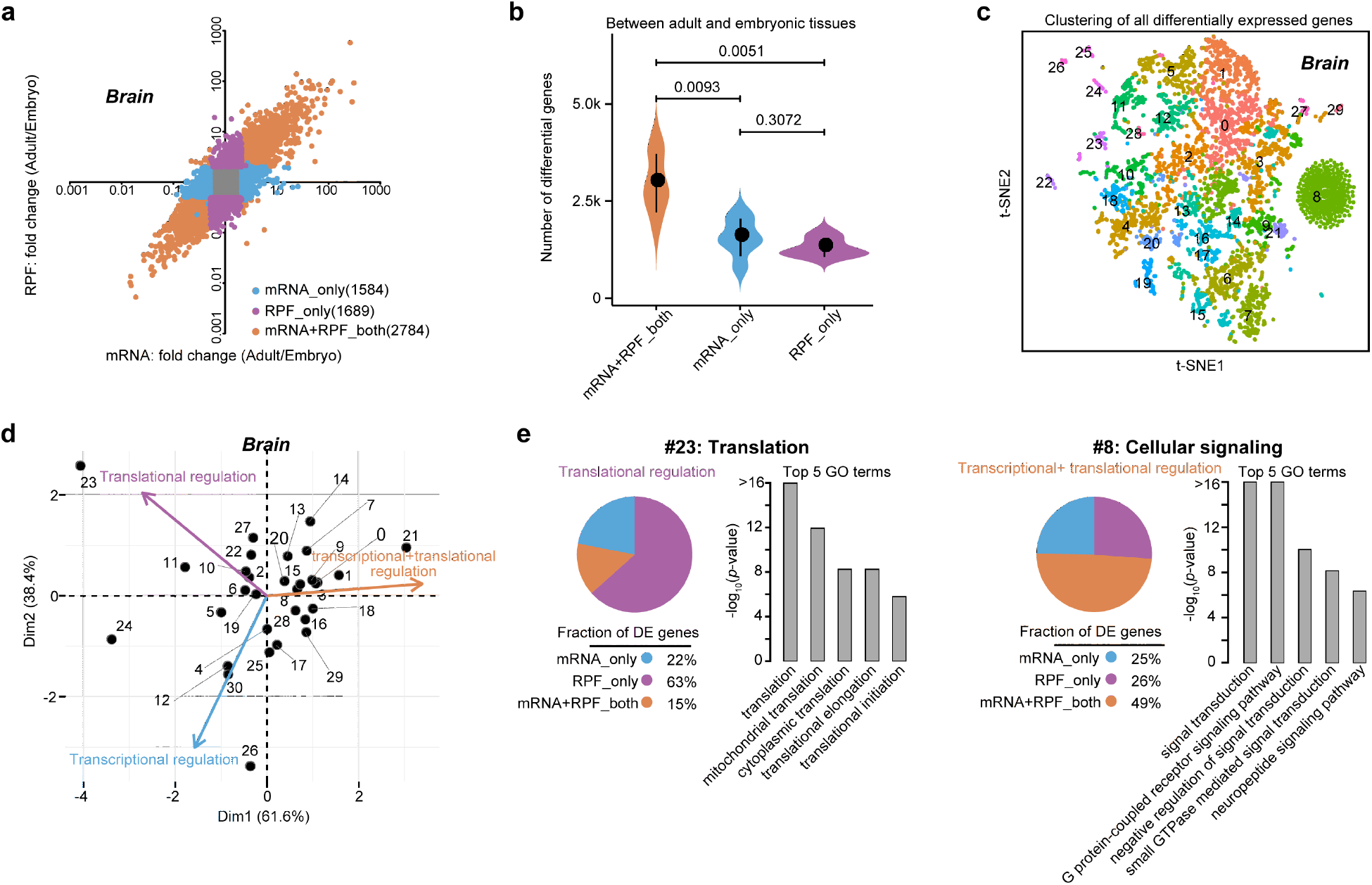
Regulatory effects of transcription and translation contribute to spatiotemporal divergences in gene expression. (**a**) Differential transcription and differential translation analyses of the same tissue among different developmental stages. Each point represents a gene. Genes with distinct patterns of differential expression are represented by different colors. (**b**) Numbers of developmental stage-specific genes exhibiting distinct patterns of differential expression. The analysis showed that the majority of differential genes were common to the transcriptome and translatome. The between-group differences were compared using a Wilcoxon rank sum test, and the *P*-values are shown. (**c**) t-SNE plots displaying the graph-based clustering results of all differentially transcribed and translated genes in the same tissue between different developmental stages. Different gene clusters are represented by different colors. (**d**) Scatter plots of the principal component analysis results showing the manifestations of individual gene clusters within the global regulatory programs. Each numbered point represents a gene cluster, and its position along each axis indicates the relative contribution of transcriptional and translational regulation to the overall differential patterns. **(e)** Examples of co-regulated gene clusters. The pie chart shows the relative fraction of differentially expressed genes with distinct differential patterns. The top-ranked GO terms and corresponding *P*-values are given. The full list of enriched GO terms for each gene cluster can be found in **Table S4**. The brain is shown in **a** and **c**-**e**, and the other tissues are shown in **Fig. S3-4**.

### Translational efficiency contributing to spatiotemporal regulation of gene expression

Translational regulation can be achieved by modulating the translational efficiency (TE), and we thus sought to quantify the spatial and temporal changes in TEs during tissue development. By comparing the TE distributions among the six tissues at the different developmental stages, we observed significant differences among the tissues (**Fig. 4a-b**). Even for the genes shared by all six tissues, we still observed substantial TE variations among the tissues (**Fig. S5a**), which illustrated that TEs are differentially used by each tissue. In addition to spatial differences, temporal differences were observed. Strikingly, all the tissues showed significantly enhanced TEs at the adult stage compared with the embryonic stage (**Fig. 4c**). Moreover, compared with the embryonic tissues, the adult tissues exhibited relatively wider ranges of TE (**Fig. 4d** and **Fig. S5b**). For instance, the TE range, which was defined as the ratio of the 97.5% to the 2.5% quantile of the TEs, spanned up to 153-fold in the adult liver, whereas the TE range in the embryonic liver spanned only 18-fold. A broad TE range might offer higher flexibility to the translational regulation of gene expression. Notably, we observed a considerably narrow spread of TEs versus transcriptional abundances for all the tissues (**Fig. 4e** and **Fig. S5c**), which indicated that the effects exerted by transcriptional control on gene expression outputs were diluted by modulating the TE. This finding might explain the effect size of the inter-tissue divergence in the translatome becoming significantly smaller than that of the transcriptome, as described in the previous section.

**Figure 4.**
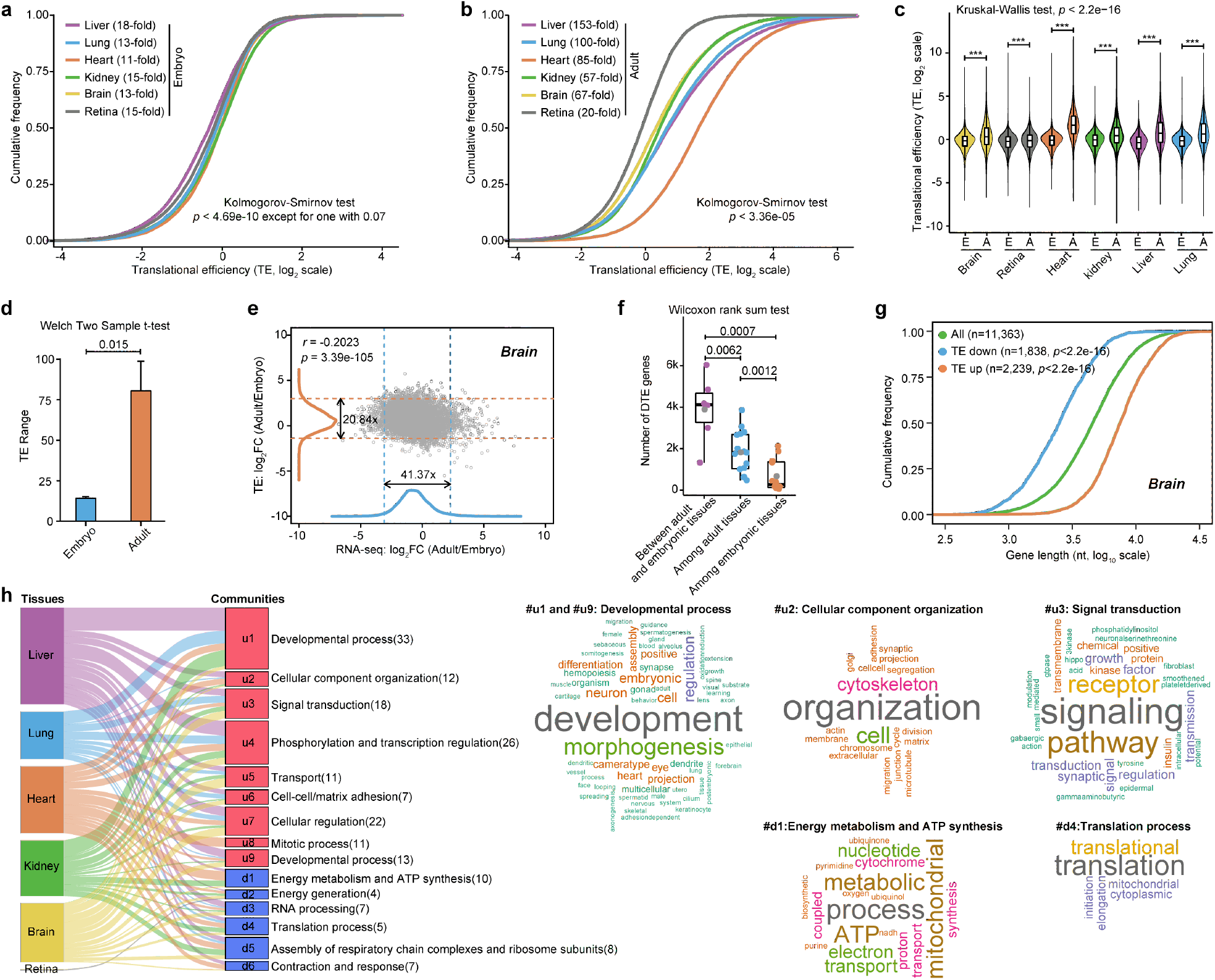
Analysis of translational efficiencies in embryonic and adult tissues. (**a**) and (**b**) Cumulative distribution of TEs of protein-coding genes expressed in embryonic and adult tissues, respectively. The between-tissue differences were compared using the Kolmogorov-Smirnov test, and all *P*-values were statistically significant except for one (*P*=0.07), which indicated a marginally significant difference between the embryonic lung and brain tissues. The TE range calculated using the formula shown below was obtained for each tissue. (**c**) Global TE differences in each tissue between the embryonic and adult stages. ***: *P*-value<0.001. The overall significance level was obtained using the Kruskal-Wallis test. (**d**) Comparison of TE ranges between embryonic and adult tissues. The differences in the TE range were compared using the Welch two-sample t-test. (**e**) Scatter plot of the adult-to-embryo ratio of transcriptional abundance versus TEs for all expressed protein-coding genes. The corresponding density curves are plotted on the margins. The dotted lines of the same colors represent the 2.5 and 97.5 percentiles of each variable, and the corresponding fold-change range is indicated. The coefficients and *P*-values for the variables in a linear regression model are presented in the left upper corner. (**f**) Comparison between the numbers of DTE genes. The between-group differences were compared using a Welch two-sample t-test, and the *P*-values are shown. (**g**) Length-dependent differential TE changes in each tissue between the adult and embryonic stages. The differentially upregulated and downregulated TE genes in adult tissue are represented by orange and blue colors, respectively. (**h**) Sankey diagram linking the functional assignments of DTE genes in each tissue between the adult and embryonic stages. The red (u1-u9) and blue boxes (d1-d6) on the right represent upregulated and downregulated GO terms, respectively. Examples of functional clusters are visualized using a WordCloud chart. The brain is shown in **e** and **g**, and the other tissues are shown in **Fig. S5**.

Moreover, thousands of differential TE (DTE) genes were detected between the two developmental stages for each tissue. Notably, the number of DTE genes between the two stages was significantly higher than the number of DTE genes among the tissues at the same stage (**Fig. 4f**), which suggested that controlling the TE is frequently used as a means of translational regulation during development. Interestingly, differential TE upregulation was found to preferentially occur in genes with longer lengths, whereas differential TE downregulation preferentially occurred in genes with shorter lengths, which were present at significantly higher levels in all tissues during development (**Fig. 4g** and **Fig. S5d**). The GO enrichment analysis showed that upregulated DTE genes were significantly enriched in biological terms associated with tissue morphogenesis, organization and architecture as well as cell-cell adhesion, transport of substances and signal transduction (**Fig. 4h** and **Table S5**), which might contribute to the enhancement of tissue maturation and increases in tissue functionality during development. In contrast, downregulated DTE genes were significantly enriched in biological terms involved in energy metabolism and ATP synthesis, such as ‘ATP metabolic process’, ‘ATP biosynthetic process’, and ‘mitochondrial electron transport’. The observed downregulation might reflect quantitative differences in the oxidative rates of mitochondria between embryonic and adult tissues. Along with these functions, translation-related functions such as ‘ribosome biogenesis/assembly’, ‘translational initiation/elongation’, and ‘cytoplasmic translation’ were also significantly downregulated in multiple adult tissues, reflecting a reduced state of protein biosynthesis in the adult tissues. Taken together, these results demonstrate that TE undergoes dynamic spatiotemporal changes that induce enhancement in tissue specificity during development.

### Regulation of spatiotemporal gene translation by uORFs

Accumulating evidence demonstrates that regulatory elements encoded by 5’UTRs play an important role in the translational regulation of gene expression. Thus, we sought to discover actively translated uORFs in each tissue during development. We detected an average of 2,049 uORFs per embryonic tissue and 1,368 uORFs per adult tissue and found large variations among the tissues ranging from 833 to 2,783 (**Fig. 5a** and see **Methods**). Significant differences in the number of uORFs and their associated genes were further found between the embryonic and adult tissues, illustrating the developmental shifts in uORF usage (*P*=0.0395 and 0.0335, respectively, one-tailed Welch two-sample t-test). In general, these uORFs tended to be highly tissue- and developmental stage-specific, as demonstrated by the finding that less than 20% of uORFs were detected in more than three tissues (**Fig. 5b**) and the low overlap of uORFs in the same tissue between two developmental stages (average Jaccard index of 0.23; **Fig. 5c**).

**Figure 5.**
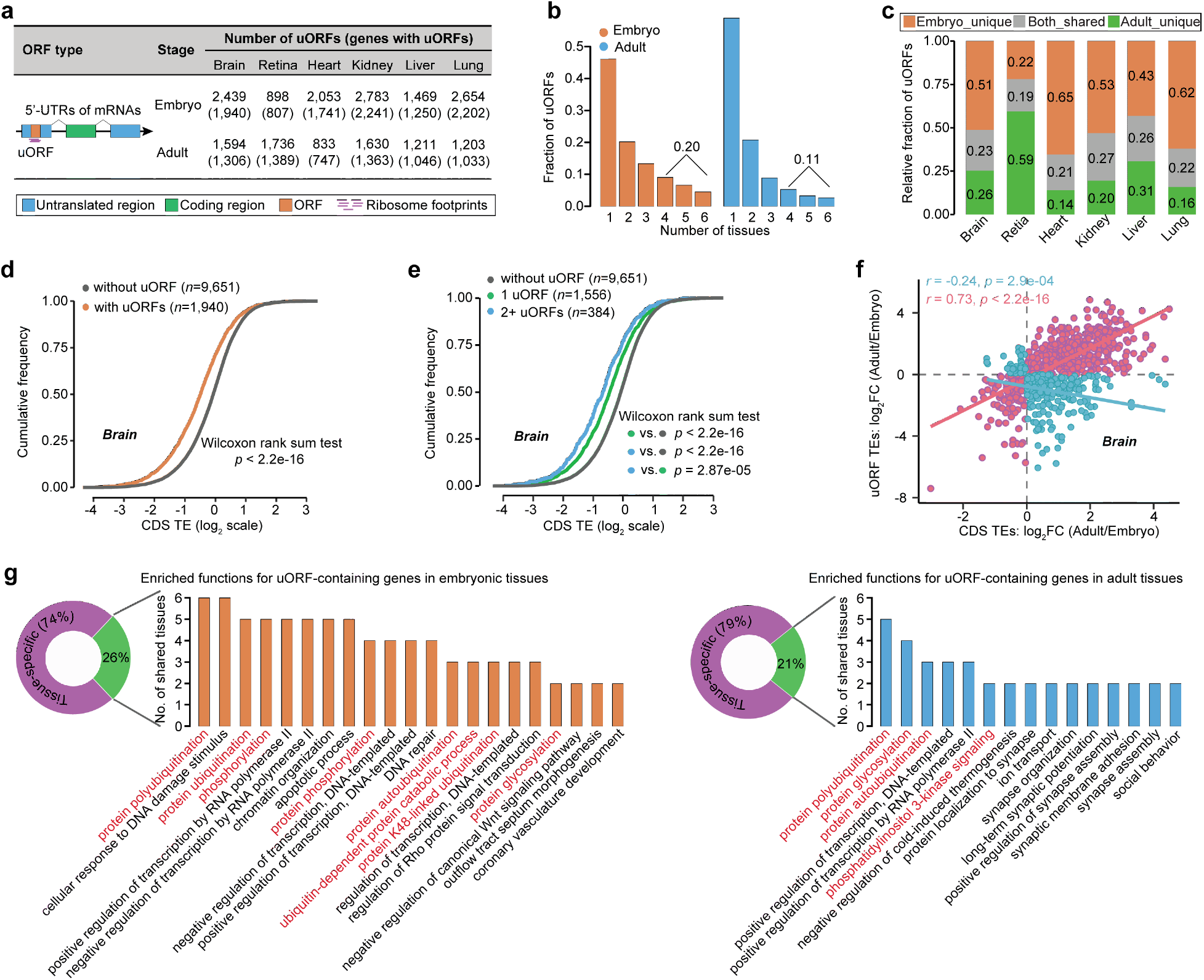
uORF-mediated translational regulation. (**a**) Numbers of uORFs and uORF-containing genes detected in each tissue. (**b**) Distribution of uORFs in embryonic and adult tissues. (**c**) Relative fraction of uORFs between different developmental stages in the same tissue. (**d**) Cumulative distribution of CDS TEs in uORF-containing genes versus those lacking uORFs. (**e**) Cumulative distribution of CDS TEs in the genes grouped by their number of uORFs. The number of uORFs is associated with a reduction in CDS TEs. (**f**) Relationship between uORF TEs and downstream CDS TEs. Each point represents an uORF. Positive and negative correlations are represented by different colors. (**g**) Representation of GO terms enriched in uORF-containing genes in embryonic and adult tissues. The full list of enriched GO terms obtained for each tissue is included in **Table S6**. The brain is shown in **d**-**f**, and the other tissues are shown in **Fig. S6**.

Given the widespread prevalence and spatiotemporal patterns of uORFs, we then explored uORF-mediated translational regulation. We observed that genes with uORFs in their leader sequences exhibited significantly reduced downstream translation, which confirmed that uORFs clearly repressed downstream translation (Johnstone et al. 2016) (**Fig. 5d** and **Fig. S6a**). This repressive effect was significantly correlated with the number of uORFs: a higher number of uORFs in a gene was associated with a stronger uORF-mediated repression of downstream translation (**Fig. 5e** and **Fig. S6b**). However, the regulatory relationships between uORFs and primary ORFs were not always conserved, particularly in adult tissues (**Fig. S6c-d**). This finding might be explained by the repressive capacity of uORFs on downstream translation being bypassed by leaky scanning or translational reinitiation (Young and Wek 2016; Zhang et al. 2019). We observed that many uORFs were positively correlated with their downstream translation, including *Eif5* and *Eef1b2*, although some uORFs were negatively correlated with their downstream translation (**Fig. 5f** and **Fig. S6e**), which illustrated the complexities of uORF-mediated regulation of gene-specific translation.

The GO enrichment analysis showed that uORF-containing genes were involved in many biological processes, some of which are directly linked to the physiological functions of each tissue type (**Table S6**). For instance, the uORFs in the embryonic brain exhibited significant enrichment in ‘nervous system development’, ‘brain development’, and ‘hippocampus development’ (*P*=3.59e-08, 3.27e-06 and 1.62e-05, respectively, hypergeometric test), and the uORFs in the adult retina exhibited significant enrichment in ‘retina development in camera-type eye’, ‘photoreceptor cell maintenance’, and ‘visual perception’ (*P*=5.98e-06, 1.58e-05 and 8.74e-05, respectively, hypergeometric test). In addition to these specific functions, many general functions, particularly those involved in post-translational modifications, such as ‘protein ubiquitination’, ‘protein phosphorylation’, and ‘protein glycosylation’ (**Fig. 5g**), were frequently observed in both embryonic and adult tissues, which suggests that uORFs are regularly used in protein modification processes to tune proteomic diversity.

Taken together, these results demonstrate that uORF-mediated translational regulation represents an important layer for the spatiotemporal control of gene expression and highlight the regulatory role and physiological importance of this control during tissue development.

### Translation of long noncoding RNA genes

In addition to protein-coding genes, the coding potentials of long noncoding RNAs (lncRNAs) have been less well studied. Taking advantage of our generated datasets, we predicted a total of 2,176 actively translated smORFs, with a median length of 50 codons (**Fig. 6a-b**, **Table S7** and **Methods)**. Notably, 92% of these smORFs were derived from annotated long intergenic noncoding RNAs (lincRNAs, 814), antisense transcripts (607) and processed transcripts (578) (**Fig. 6c**), including previously well-known translated lncRNAs, such as *Neat1* and *Dancr* (van Heesch et al. 2019). We then used publicly available Ribo-seq datasets to validate our predicted smORFs (see **Methods**), and a re-analysis of these public dataset confirmed 723 of the 2,176 smORFs. Using mass spectrometry-based proteomic data, we obtained direct peptide evidence for micropeptides translated from 203 of the 2,176 smORFs (**Fig. 6d**). In addition, *in vitro* translation experiments were performed for 10 randomly chosen smORFs, and three of these, namely, *Lsmem2*, *RP23-83I13.10*, and *RP23-52N2.1*, were found to generate translation products (**Fig. 6e**). Given the spatiotemporal transcriptional dynamics of lncRNAs (Deveson et al. 2017), we subsequently examined the translational dynamics of smORFs during tissue development. We found that most of the smORFs were translated with high spatiotemporal specificity; specifically, 55% and 70% of the smORFs were translated in only one tissue at the embryonic and adult stages, respectively (**Fig. 6f**), and more than 68% of the smORFs were translated at one stage (**Fig. 6g**). Highly specific spatiotemporal patterns of smORF translation suggest their potential functional importance for development.

**Figure 6.**
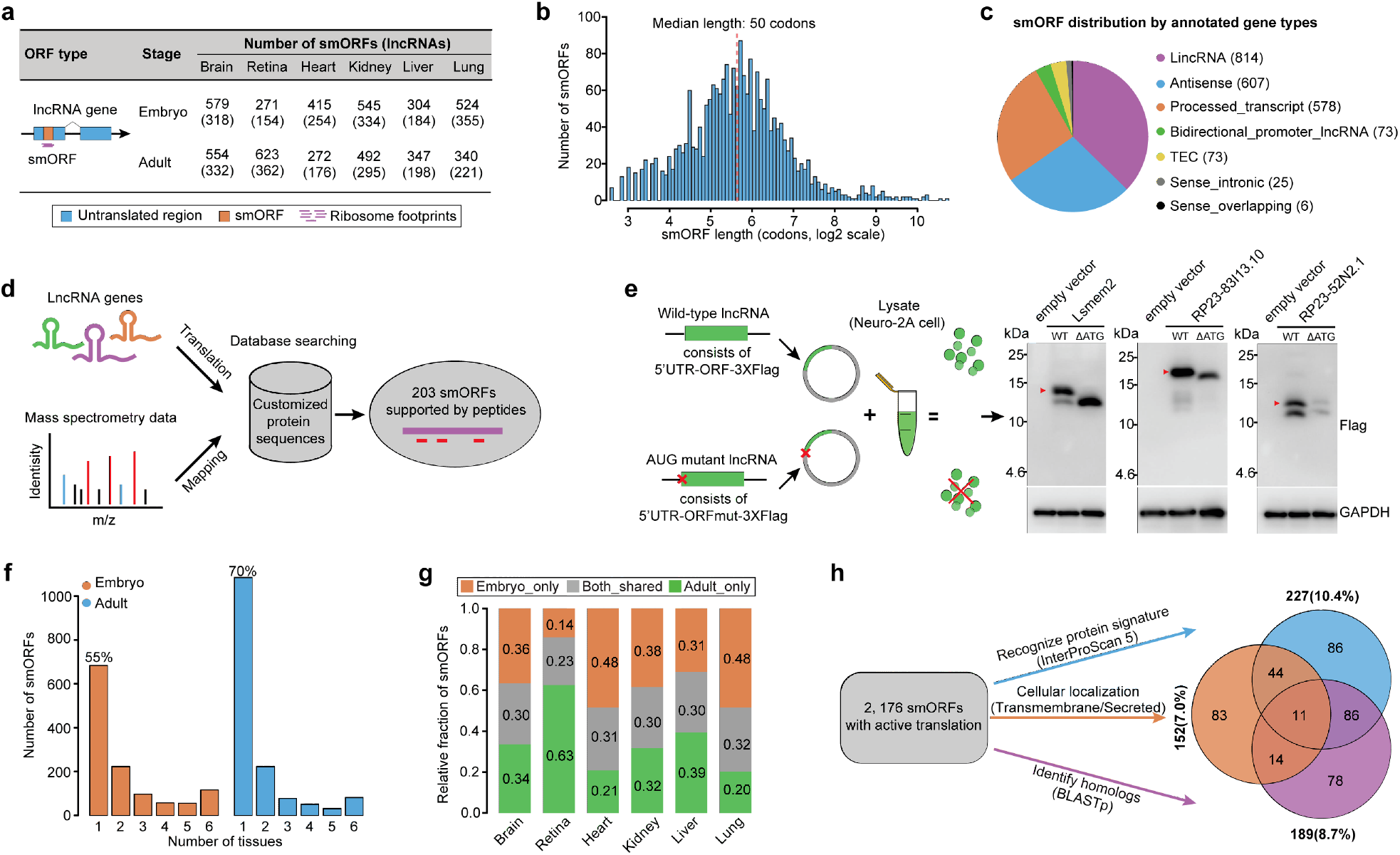
lncRNA translation. (**a**) Number of actively translated smORFs detected in each tissue. (**b**) Length distribution of the identified smORFs, with a median size of 50 codons. (**c**) smORF distribution by annotated gene types. The largest fraction was obtained with lincRNAs. (**d**) *In vivo* assessment of smORF translation using mass spectrometry-based proteomic data. (**e**) *In vivo* experiments for validating smORF translation. Molecular weights of micropeptides are indicated in kilodaltons (kDa). (**f**) and (**g**) Translational dynamics of smORFs exhibiting spatiotemporal specificity. (**h**) Functional characterization of smORF-encoded peptides (SEPs). Venn chart showing the overlap among numbers of SEPs annotated with different tools.

We subsequently performed a systematic *de novo* annotation of the smORF-encoded peptides (SEPs) to understand their functional potentials (**Fig. 6h**; see **Methods**). The sequences of these putative SEPs were first searched against a set of protein signature databases. Of the 2,176 SEPs, 227 (~10.4%) were predicted to have identifiable features found in known proteins, such as conserved domains, functional sites, and protein family information, and notably, some of these were assigned to certain molecular functions, such as a 31-amino-acid (aa) SEP from *RP23-272H13.2* involved in ‘DNA-binding transcription factor activity’. Because peptide signals mediate a variety of cell-to-cell communications crucial for growth and development, we subsequently performed transmembrane topology and signal peptide predictions to search for SEPs that can potentially act as mediators of cellular communications. A total of 152 (~7.0%) SEPs were predicted to be transmembrane and/or secreted, and these included 84 that were solely transmembrane, 55 that were solely secreted, and 13 that were both transmembrane and secreted. We then performed sequence similarity searches of all the SEPs against known small proteins with less than 100 aa annotated in the mouse reference genome. A total of 189 (~8.7%) SEPs were identified to have recognizable homologs of known small proteins. Intriguingly, all these annotated SEPs (402) strongly tended to be enriched in multiple tissues (*P*<2.2e-16, Fisher’s exact test), which suggested their potential important physiological roles in developing tissues. Taken together, these results indicated that many SEPs exhibited remarkable functional relevance. Moreover, the majority did not have any known structural, localization or functional properties, which imposes a great challenge on the functional annotation of the smORFs.

## Discussion

Translational regulation plays critical roles in tissue functionality and development. Our translatome analysis provided many novel insights into the translational regulation that modulates the spatiotemporal dynamics of gene expression and physiological functions during development. We quantified the spatiotemporal divergences of global gene expression in developing tissues. The functional characterization of genes and pathways underlying the divergences in gene expression enhanced our understanding of the molecular basis of tissue physiology. We found that ubiquitously translated genes are involved in many enriched pathways that are important for the maintenance of basic cellular functions. Notably, translational profiling will enable a better definition of ‘housekeeping genes’, as supported by the observation that approximately 14% of ubiquitously transcribed genes were not included in the set of ubiquitously translated genes. This change is likely subject to disallowance of the regulation during the process transition from transcription to translation. Tissue-enriched translated genes exhibited higher levels of translation compared with ubiquitously translated genes, which suggests that translational regulation plays an important role in fulfilling tissue function. This finding could be further confirmed by the observation of the substantial use of TE regulation among different tissues, including spatiotemporal TE changes and uORF usages. Many uORFs provide functionally important repression of downstream primary CDS in a dose-dependent manner, consistent with some previous reports (Bazzini et al. 2014; Johnstone et al. 2016). However, some uORFs exert different regulatory effects on the translation of primary CDS, particularly in adult tissues, demonstrating the complexities of uORF-mediated translational regulation (Zhang et al. 2019). These findings illuminate the importance of translational regulation in shaping the gene repertoires expressed in developing mouse tissues.

The identification of actively translated smORFs in annotated lncRNAs broadens our understanding of the protein-coding potential of the genome. We revealed a large number of translated smORFs that exhibit spatiotemporal patterns. However, as demonstrated in our previous study (Wang et al. 2017), smORFs are also obviously different from annotated protein-coding genes and exhibit markedly distinguishing properties, including expression, structural, sequence, evolutionary, and functional features, which underpin the mysteries surrounding the biology of lncRNAs. Although our analysis revealed pervasive actively translated smORFs in different tissues and stages, translatable smORFs are not necessarily detectable smORFs. Indeed, only a portion of our identified smORFs were validated by different strategies, including publicly available Ribo-seq data, *in vivo* micropeptide detection by mass spectrometry, and *in vitro* translation experiments. One possibility is that some SEPs likely escape detection due to extremely low expression abundance (Wang et al. 2019), and another possibility is that some SEPs are likely degraded during proofreading through nonsense-mediated decay (Makarewich and Olson 2017). In addition, detectable SEPs are also not necessarily functional SEPs. To understand the functional potentials of SEPs, we used integrative annotation approaches and provided the functional relevance of 402 SEPs. Nevertheless, understanding the full repertoire of translatable smORFs and their functions remains challenging.

In summary, our analyses presented here facilitate a better understanding of how tissue-specific and developmental stage-specific phenotypes are achieved through the precise spatiotemporal control of translational regulation and can also be used as a useful resource for translatome research.

## Materials and Methods

### Tissue collection

Wild-type C57BL/6 mice were purchased from the Guangdong Medical Experimental Animal Center (Guangdong, China; License No: SCXK (YUE) 2018-0002). Brain, heart, kidney, liver, lung, and retinal tissues were harvested separately from embryonic (E15.5) and adult (P42) C57BL/6 mice and immediately snap frozen in liquid nitrogen. All experimental procedures were approved by the Animal Ethics Committee of the Zhongshan Ophthalmic Center, Sun Yat-sen University (Guangzhou, China; License No: SYXK (YUE) 2018-0189), in accordance with institutional animal welfare guidelines and the Animal Protection Law of China.

### Library preparation and sequencing

Frozen tissue samples were lysed using 1 ml of mammalian lysis buffer (200 μl of 5x Mammalian Polysome Buffer, 100 μl of 10% Triton X-100, 10 μl of DTT (100 mM), 10 μl of DNase I (1 U/μl), 2 μl of cycloheximide (50 mg/ml), 10 μl of 10% NP-40, and 668 μl of nuclease-free water). After incubation for 20 minutes on ice, the lysates were cleared by centrifugation at 10,000xg and 4°C for 3 minutes. For each tissue and replicate sample, the lysate was divided into 300-μl and 100-μl aliquots. For the 300-μl aliquots of clarified lysates, 5 units of ARTseq Nuclease was added to each A_260_ lysate, and the mixtures were incubated for 45 minutes at room temperature. Nuclease digestion was stopped by the addition of 15 μl of SUPERase·In RNase Inhibitor (Ambion). Subsequently, the lysates were applied to Sephacryl S-400 HR spin columns (GE Healthcare Life Sciences), and ribosome-protected fragments were purified using the Zymo RNA Clean & Concentrator-25 kit (Zymo Research). Ribosomal RNA was depleted using the Ribo-Zero magnetic kit (Epicentre). Sequencing libraries of ribosome-protected fragments were generated using the ARTseq™ Ribosome Profiling Kit (Epicentre, RPHMR12126) according to the manufacturer’s instructions. From the 100-μl aliquots of clarified lysates, poly(A)+ RNAs were extracted and purified, and sequencing libraries of poly(A)+ RNAs were then generated using the VAHTS™ mRNA-seq v2 Library Prep Kit from Illumina (Vazyme Biotech, NR601-01) according to the manufacturer’s instructions. The resulting 48 barcoded libraries were pooled and sequenced using an Illumina HiSeq 2500 instrument in the single-end mode.

### Sequencing data preprocessing

The raw sequence reads were demultiplexed using CASAVA (version 1.8.2), and the 3’-end adapter was clipped using Cutadapt (version 1.8.1) (with the parameters ‘-a AGATCGGAAGAGCACACGTCTGAACTCCAGTCA-match-read-wildcards-m 6’). Low-quality sequences were trimmed using Sickle (version 1.33) (with the parameters ‘-q 20’). The trimmed reads were filtered by length based on the ranges [25, 34] for ribosome footprints and [20, 50] for mRNA. The retained reads that mapped to reference mouse rRNAs or tRNAs were then removed, and the remaining reads were aligned to the mouse reference genome (downloaded from GENCODE, Release M18: GRCm38.p6) using Tophat2 (version 2.0.14) (Kim et al. 2013) with the following command: ‘tophat2 -g 20 -N 2 --transcriptome-index [index_file] -G [gtf_file] [fastq_file] -o [output_directory]’. Only those uniquely mapped reads were extracted for gene expression determination. The total number of aligned count reads per gene was obtained using the Subread R package-featureCounts (version 1.6.2) (Liao et al. 2019) and further converted to transcripts per kilobase million (TPMs).

### Defining expressed genes

To determine putative genes expressed at levels that are significantly higher than the background levels, a half-Gaussian distribution of expression values (log2(TPM)) for each dataset was fitted through kernel density estimation using the ks R package (Duong 2018). The half-Gaussian was then mirrored to a full Gaussian distribution. A two-fold standard deviation below the mean of the distribution was chosen as the minimum threshold for gene expression. Different threshold values, which were defined in a sample-specific manner, were used to filter low-abundance genes. Genes below the threshold in any one replicate of each tissue were filtered out (that is, not included) in the subsequent analyses. Additionally, the translated genes were further required to contain actively translated ORFs (see below). It should be noted that this filtering process reduced the noise in gene expression and thus provided high-confidence quantitation for gene expression, as demonstrated by the observation that the low-abundance genes that did not pass the threshold exhibited substantially weaker correlations (mean Pearson correlation coefficients r=0.59 and 0.50 for the embryonic and adult tissues, respectively) between two replicates of each tissue.

### Gene classification

The development of different omics-based analyses have allowed the classification of protein-coding genes with regard to tissue-restricted expression. In line with this, we took advantage of the algorithm provided in the Human Protein Atlas (Kim et al. 2014) to define tissue-specific genes that can be grouped as follows: (1) ‘tissue-enriched’ genes, defined as genes showing at least five-fold higher expression in one tissue compared with all other tissues; (2) ‘group-enriched’ genes, defined as genes showing at least five-fold higher expression in a group of tissues (2-5) compared with all other tissues; (3) ‘expressed-in-all’ genes, defined as genes expressed in all analyzed tissues; (4) ‘not detected’ genes, defined as genes not present in any of the analyzed tissues; and (5) ‘mixed’ genes, defined as genes not belonging to any part of the other categories. The detailed RNA-seq- and Ribo-seq-based classifications of all mouse protein-coding genes conducted using the TissueEnrich R package (Jain and Tuteja 2019) are included in **Table S2**.

### Detection of actively translated ORFs

Canonical and noncanonical ORF detection was performed using Ribo-TISH (version 0.2.1) (Zhang et al. 2017) with the longest strategy under the default threshold setting, which uses a frame test based on the nonparametric Wilcoxon rank-sum test to determine the significance of 3-nt periodicity in the P-site signals along an ORF. Notably, to increase the statistical power of the ORF identification, the aligned BAM files for two replicates per tissue were merged together with ‘samtools merge’ (version 1.6), and only those uniquely mapped reads were used in the Ribo-TISH analysis. The final set of actively translated ORFs with an AUG-start codon followed by an in-frame stop codon in annotated protein-coding transcripts was stringently filtered based on the requirement of a minimum length of 18 nucleotides and the expression of the ORF-containing gene at an above-background level. uORFs were defined as ORFs originating from the 5’UTRs of annotated protein-coding genes, and smORFs were defined as ORFs originating from annotated long noncoding genes.

### Calculation of gene expression divergence

The divergence of expression profiles between a pair of tissues was measured based on the Euclidean distance (Glazko and Mushegian 2010), which was defined as follows:

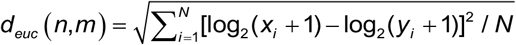

where *x*_*i*_ and *y*_*i*_ represent the expression values of gene *i* in tissues *n* and *m*, respectively, and *N* represents the number of protein-coding genes used in the comparison of global gene expression patterns (here is 17,211). Notably, this metric was used for all expression divergence calculations presented in this study, and the general patterns of divergence between tissues observed using this metric were further confirmed through a tissue specificity analysis (see below).

### Tissue specificity analysis

The tissue specificity of each gene was determined using the τ index (Yanai et al. 2004) as follows:

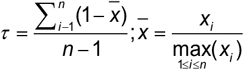

where *n* represents the number of tissue types and *x*_*i*_ represents the average TPM value of the gene in tissue type *i*. This index varies on a scale from 0 to 1, where 0 indicates ubiquity and 1 indicates specificity.

### Gene ontology (GO)-based enrichment analysis

All GO term annotations for Mouse Genome Informatics (MGI) were extracted from the ‘mgi.gaf.gz’ file that was downloaded from the Gene Ontology website (http://current.geneontology.org/products/pages/downloads.html). After assigning all genes to GO terms, only those GO terms for biological processes containing at least five genes were retained for functional enrichment analysis. In total, 17,596 genes assigned to 4,896 GO terms were included in this analysis. The hypergeometric distribution was further used to determine whether a GO term was overrepresented in a given gene set. After multiple testing correction using the Benjamini-Hochberg (BH) approach, those GO terms with a false discovery rate (FDR) below 5% were determined to be statistically significant.

### Differential expression analysis

To allow proper comparisons among the RNA-seq and Ribo-seq data, raw read counts obtained at the exon level using featureCounts were combined together and normalized against the reference to yield a pool-based size factor, and the resulting data were used for differential expression analysis with the DESeq2 R package (Love et al. 2014). A gene was considered to be significantly differentially transcribed or translated if it met the following criteria: (1) the FDR was controlled at the 5% level and (2) the absolute fold-change (FC) threshold was set to the most typical cutoff value of 2 (FC>2 and FC<1/2). After characterizing concordant and discordant changes in transcription and translation, we defined three distinct patterns of differential genes: ‘mRNA+RPF_both’, which indicated concordant differential expression in both transcription and translation; ‘mRNA_only’, which indicated differential expression in transcription but not in translation; and ‘RPF_only’, which indicated differential expression in translation but not in transcription.

### Cluster assignment of co-regulated genes

The pairwise semantic similarities of all differentially transcribed and translated genes across each pairwise comparison were first calculated using the GOSemSim R package (Yu et al. 2010) to define gene clusters that were expected to exhibit co-regulatory functional arrangements. The similarity matrix composed of pairwise semantic similarities of genes was then used as the input for a graph-based clustering analysis. A t-SNE plot from the Seurat R package (Satija et al. 2015) was used to visualize the output of the clustering analysis.

### Principal component analysis

A PCA was performed to dissect the relative contribution of gene regulation to the overall differential pattern of each cluster. For each cluster of co-regulated genes, we calculated the fractions of previously defined genes with three different differential patterns that were used as the input for the PCA. The prcomp and fviz_pca_biplot functions from the factoextra R package (Kassambara and Mundt 2017) were used for the PCA and visualizing the output of the PCA, respectively.

### Estimation of the translational efficiency

The TE for each protein-coding gene was estimated as the ratio of the normalized values (TPM) of Ribo-seq to RNA-seq reads in the annotated CDS region. A similar definition was used for estimating the TE of uORFs. Briefly, following normalization to the TPM value, the sequenced reads uniquely mapped to each of the uORFs were counted with featureCounts. The expression measurements per uORF were then averaged across two replicates. The TE of each uORF was calculated as the ratio of the average TPM values of the Ribo-seq to RNA-seq reads.

### Analysis of differential translational efficiency

The changes in the TE of a gene between and within different adult tissues and their corresponding embryonic tissues were assessed using the DESeq2 R package (Love et al. 2014) with a threshold of 0.05 to control the FDR and an absolute FC > 2. A table of raw read counts within the whole CDS regions obtained using featureCounts was used as the input for this analysis.

### Analysis of publicly available ribosome profiling data

Twenty-five publicly available Ribo-seq datasets (Accession in **Table S7**) were downloaded from the GEO database, and only the matched samples from our analyzed tissue types were retrieved for subsequent analysis. The raw data files were preprocessed, and all smORFs were predicted using the same procedures as those used in our previous analyses. This set of smORFs was used to determine if any of our predicted smORFs exhibited active translation in the publicly available datasets.

### Analysis of publicly available mass spectrometry-based proteomic data

Two publicly available proteomic datasets were retrieved from the PRIDE database (Accession number: PXD009909) and other public resources from the PRIDE database and other public resources (https://phosphomouse.hms.harvard.edu/data/). The first included samples from five of our analyzed tissue types, and the second included samples of another tissue type. The raw data files were analyzed using MaxQuant software (version 1.6.10.43) (Cox and Mann 2008) against a custom-made database, which combined all mouse sequences from UniProt/Swiss-Prot (MOUSE.2019-06) with sequences derived from translatable lncRNAs, based on the target-decoy strategy (Reverse) with the standard search parameters with the following exceptions: (1) the peptide-level FDR was set to 5%, and the protein-level FDR was excluded; (2) the minimal peptide length was set to six amino acids; and (3) a maximum of two missed cleavages was allowed. Finally, a total of 203 smORF-encoded peptides (SEPs) were evidenced by at least one unique peptide.

### *In vitro* translation experiments

#### Plasmid constructs

To generate 3xFlag fusion protein constructs, smORF sequences with endogenous pseudo 5’UTRs (defined as the region upstream of the smORF start codon) were amplified by RT-PCR and then cloned into the pcDNA3.1-3xFlag vector, which is a homemade plasmid from pcDNA3.1(+) (Invitrogen). A mutation construct (5’UTR-ORFmut-3xFlag) in which the smORF start codon was mutated to ATT was generated using a Mut Express II Fast Mutagenesis Kit V2 (Vazyme). The wild-type and mutant plasmids were verified by Sanger sequencing. The primers used in this study are listed in **Table S7**.

#### In vitro translation

Both wild-type and mutant plasmids were transfected into Neuro-2A cells using Lipofectamine 3000 reagent (Invitrogen), and 48 hours later, the cells were harvested and resuspended in RIPA (Beyotime) with protease inhibitor cocktail (Roche). The cellular lysates were denatured at 85°C for 5 minutes and then separated on 16.5% Tricine gels for 1 hour at 30 V and then for 4 hours at 100 V. The proteins were then electroblotted onto a polyvinylidene fluoride (PVDF) membrane (Millipore), and the PVDF membranes were then blocked in 5% nonfat dry milk in TBST for 1 hour. Western blotting was performed using anti-Flag (1:1000) (Sigma) or anti-GAPDH (1:5000) (Proteintech) primary antibodies, and the membranes were incubated with the secondary antibodies conjugated to horseradish peroxidase (anti-mouse from CST, 1:10000) for 1 hour. The Western blotting signals were developed using Immobilon Western Chemiluminescent HRP Substrate (Millipore) and imaged with ChemiDoc™ Imaging Systems (Bio rad).

### *De novo* functional annotation of smORF-encoded peptides

#### Protein signature recognition

Each of the putative smORF-encoded peptides (SEPs) was queried against a default set of protein signature databases, including CATH-Gene3D, CDD, PANTHER, Pfam, PIRSF, PRINTS, and SMART, through a local InterProScan 5 search (version 5.35-74.0) (Jones et al. 2014) with a hit e-value threshold of 0.0001.

#### Cellular localization prediction

Transmembrane and secreted SEPs were predicted using the web applications TMHMM 2 (http://www.cbs.dtu.dk/services/TMHMM/) and SignalP-5.0 (http://www.cbs.dtu.dk/services/SignalP/) with the default parameters. The predictions provided the most likely location and orientation of the transmembrane helices in the sequence as well as the presence of signal peptides and the location of their cleavage sites in the proteins.

#### Protein homology detection

All mouse protein-coding transcript translation sequences were downloaded from GENCODE (Release M18: GRCm38.p6), and the sequences of proteins composed of less than 100 aa were further retrieved as the set of known small proteins. Each SEP was then queried against this set of known small proteins using BLASTp software (version 2.7.1+) (Altschul et al. 1990) with a hit e-value threshold of 0.05.

### Data access

All raw Ribo-seq and RNA-seq data generated in this study have been submitted to the NCBI Gene Expression Omnibus (GEO; https://www.ncbi.nlm.nih.gov/geo/) under accession number GSE94982.

## Acknowledgments

This work was supported by the National Natural Science Foundation of China [31871302, Z.X.], the Joint Research Fund for Overseas Natural Science of China [31829002, Z.X.] and the Center for Precision Medicine at Sun Yat-sen University.

## Author contributions

H.W.W. and Z.X. designed the study and wrote the manuscript. J.Q.Y. and N.T. performed the experiments. H.W.W., Y.W., and H.H.L. performed the bioinformatics analyses. M.Z.X. constructed the database. All named authors read and approved the final manuscript.

## Supplementary Figures and Tables

**Figure S1.** Data quality evaluation. (**a**) Distribution of uniquely mapped reads in different genomic regions in each sample. (**b**) Pearson correlation coefficients obtained from pairwise correlation analyses of all Ribo-seq and RNA-seq datasets. The correlations between biological replicates along the diagonal are highlighted with bold font. (**c**) Principal component analysis of Ribo-seq and RNA-seq data using protein-coding genes with TPM values greater than 1 in at least three samples.

**Figure S2.** Tissue specificity of gene expression measured using the Tau-index metric. The between-group differences were compared using a Wilcoxon rank sum test, and the *P*-values are shown.

**Figure S3.** Differential gene expression analysis. (**a**) Scatter plots showing differentially transcribed and translated genes in the same tissue among different developmental stages. The analysis was performed using DESeq2 with exon-level counts (false discovery rate, FDR<0.05 and |log2-fold change|>1). Each point represents a gene, and genes with distinct patterns of differential expression are represented by different colors. (**b**) Numbers of differentially transcribed and translated genes among embryonic and adult tissues, respectively. The between-group differences were compared using a Wilcoxon rank sum test, and the *P*-values are shown.

**Figure S4.** Dissection of transcriptional and translational regulation during development. (**a**) t-SNE plots displaying the graph-based clustering results of all differentially transcribed and translated genes in the same tissue among different developmental stages. The gene clusters are represented by different colors. (**b**) Scatter plots of the PCA results showing the manifestations of individual gene clusters within the global regulatory programs. Each numbered point represents a gene cluster, and its position along each axis indicates the relative contribution of its transcriptional and translational regulation to the overall differential patterns.

**Figure S5.** Analysis of the translational efficiencies in embryonic and adult tissues. (**a**) Cumulative distribution of TEs of protein-coding genes shared by all six tissues. (**b**) Comparison of TE ranges of protein-coding genes shared by all six tissues between the embryonic and adult stages. The differences in the TE ranges were compared using the Welch two-sample t-test. (**c**) Scatter plots of the adult-to-embryo ratio of the transcriptional abundance versus the TEs for all expressed protein-coding genes. The corresponding density curves are plotted on the margins. Dotted lines of the same colors represent the 2.5 and 97.5 percentiles of each variable, and the corresponding fold-change range is indicated. The coefficients and *P*-values for the variables in a linear regression model are presented in the left upper corner. (**d**) (**g**) Length-dependent differential TE changes in each tissue between the adult and embryonic stages. The differentially upregulated and downregulated TE genes in adult tissue are shown in orange and blue, respectively.

**Figure S6.** uORF-mediated translational regulation. (**a**) and (**c**) Cumulative distribution of CDS TEs in uORF-containing genes versus those lacking uORFs. (**b**) and (**d**) Cumulative distribution of CDS TEs in the genes grouped by their number of uORFs. The number of uORFs is associated with a reduction in CDS TEs. (**e**) Relationship between uORF TEs and downstream CDS TEs. Each point represents an uORF. Positive and negative correlations are represented by different colors.

**Table S1**. Summary of Ribo-seq and RNA-seq data from the six different tissues.

**Table S2**. Gene classification based on their transcriptional and translational profiles.

**Table S3**. Enriched GO terms for tissue-enriched and expressed-in-all genes.

**Table S4**. Enriched GO terms for co-regulatory gene clusters.

**Table S5**. Enriched GO terms for differential TE genes.

**Table S6**. Enriched GO terms for uORF-containing genes.

**Table S7**. Detailed information on smORFs encoded by lncRNAs.

